# Interplay of NuA4/TIP60 and PRC2 Complex Activities in Fusion Driven Endometrial Stromal Sarcoma

**DOI:** 10.64898/2026.09.04.749526

**Authors:** Deepthi Sudarshan, Charles Joly-Beauparlant, Stephanie Bianco, Catherine Lachance, Maëlys Le Goff, Nader Alerasool, Elliot Gregoire, Lara Hermann, Christophe Tav, Jean-Philippe Lambert, Marcus Q. Bernardini, Arnaud Droit, Mikko Taipale, Jacques Côté

**Affiliations:** Centre Hospitalier Universitaire (CHU) de Québec-Université Laval Research Center, Laval University Cancer Research Center, Québec City, Québec G1R 3S3, Canada; Donnelly Centre for Cellular and Biomolecular Research, Department of Molecular Genetics, University of Toronto, Toronto, Ontario M5S 3E1, Canada; Computational Biology Laboratory, CHU de Québec-Université Laval Research Center, Québec City, Québec G1V 4G2, Canada; Department of Gynecologic Oncology, Princess Margaret Cancer Center, University Health Network, Sinai Health System, Toronto, Ontario M5B 2M9, Canada; Department of Obstetrics and Gynecology, University of Toronto, Toronto, Ontario M5G 1X8, Canada

## Abstract

Low-grade endometrial stromal sarcoma (LGESS) exhibits frequent chromosomal translocations that fuse various subunits of the NuA4/TIP60 co-activator complex to subunits of the Polycomb Repressive Complex 2 (PRC2) complex. LGESS fusion proteins, such as the commonly occurring JAZF1-SUZ12, have been shown to upregulate genes through mislocalization of NuA4/TIP60 activity to Polycomb target genes. In this study, we characterized an interesting recurrent fusion protein in LGESS that fused a NuA4/TIP60 component, MBTD1, to EZHIP. EZHIP is a recently described vertebrate protein that enzymatically inhibits methyltransferase activity of the PRC2 complex and is a potent oncogene. The MBTD1-EZHIP fusion protein forms a chimeric TIP60-PRC2.1 complex and drastically reduces H3K27me3 levels at Polycomb target genes, similar to EZHIP-overexpressing cancers. However, unlike EZHIP overexpression, MBTD1-EZHIP requires mislocalization of NuA4/TIP60 activity through the MBTD1 protein to upregulate oncogenes. Despite differences in the finer molecular mechanisms, MBTD1-EZHIP and the common JAZF1-SUZ12 fusion protein upregulated similar sets of genes in cell lines, showing a convergence of oncogenic gene expression. Surprisingly, unlike in cellular models, *JAZF1-SUZ12* translocated patient samples showed not only upregulation of genes driven by increased H4K8ac and decreased H3K27me3 but also downregulation of specific genes due to accumulated H3K27me3, revealing an additional oncogenic mechanism in LGESS.

## INTRODUCTION

Sarcomas are rare connective tissue cancers that frequently harbor chromosomal translocations. Chromosomal translocation is a pathognomonic modification in such sarcomas, with few other genetic aberrations. Many chromosomal translocations in sarcomas involve chromatin-modifying proteins that alter the cellular transcriptome. One such class of sarcoma is Endometrial Stromal sarcoma (ESS), which is a rare form of uterine cancer, accounting for approximately 1% of all uterine cancer cases (Brandon-Luke et al. 2017). These sarcomas originate from stromal cells of the endometrium and resemble proliferative-phase endometrial stromal cells. The low-grade form of this cancer (LGESS) is characterized by frequent chromosomal translocations that fuse genes encoding subunits of the NuA4/TIP60 complex (EPC1/2, MBTD1, MEAF6, BRD8, JAZF1, and ING3) with genes encoding components of the Polycomb Repressive Complex 2 (PRC2) (PHF1/SUZ12/EZHIP/EZH2/EED), with the JAZF1-SUZ12 fusion being the most prevalent (Ferreira et al. 2018). These specific fusions drive these cancers, demonstrating a low mutational burden and distinct clustering of gene expression within the LGESS class. Thus, these cancers involve the fusion of protein subunits from distinct chromatin-modifying complexes, each with opposing functions in gene regulation. Specifically, the NuA4/TIP60 histone acetyltransferase (HAT) complex is involved in H4, H2A, and H2A.Z acetylation and exchange of canonical H2A with H2A.Z, leading to transcriptional activation (Doyon et al. 2004; Sapountzi and Côté 2010)(Jacquet et al. 2016; Yang et al. 2024). Conversely, Polycomb Repressive Complex 2 (PRC2) represses transcription through H3K27me3 and genome organization (Schuettengruber et al. 2017).

Studies by our group and others have revealed that the JAZF1-SUZ12 and EPC1-PHF1 fusion proteins found in LGESS assemble a chimeric complex of NuA4/TIP60 and PRC2. The primary molecular mechanism of these TIP60-PRC2 fusions is the increase in NuA4/TIP60 mediated H4 acetylation and loss of PRC2 mediated H3K27me3 at bivalent/Polycomb target genes (Sudarshan et al. 2022; Tavares et al. 2022; Alerasool et al. 2022; Piunti et al. 2019).

A recently characterized PRC2 complex-interacting protein, EZHIP/CXORF67, is recurrently fused to MBTD1, a member of the NuA4/TIP60 complex, resulting in the MBTD1-EZHIP fusion protein in LGESS (Dewaele et al. 2014). EZHIP (EZH2 Inhibitor Protein), usually expressed only in the ovaries, is overexpressed in posterior fossa type A (PFA) ependymomas, resulting in allosteric inhibition of PRC2 activity. The C-terminus of EZHIP contains a stretch of amino acids that resemble the H3K27M onco-histone mutation found in High-Grade Gliomas (HGG). The H3K27M mimic (KLP) region in EZHIP acts similarly to the H3K27M mutant and reduces bulk H3K27me3. On chromatin, H3K27me3 is restricted from spreading to large domains and remains at the H3K27me3 nucleation sites (Hübner et al. 2019; Ragazzini et al. 2019; Piunti et al. 2019; Jain et al. 2019).

A previous study reported that the MBTD1-EZHIP fusion protein interacts with NuA4/TIP60 and PRC2 complex proteins. Western blot experiments in cells expressing MBTD1-EZHIP indicated a global loss of H3K27me3 (Piunti et al. 2019), whereas studies of other TIP60-PRC2 fusions in LGESS showed no bulk loss of H2K27me3. Moreover, since EZHIP expression in non-germ cells can be strongly oncogenic, the role of NuA4/TIP60 associated with EZHIP in the MBTD1-EZHIP fusion is unclear. Furthermore, the gene expression profiles of MBTD1-EZHIP and JAZF1-SUZ12 in LGESS patient samples clustered together when compared with High-Grade ESS (HGESS) or Undifferentiated Stromal Sarcoma (USS) (Dewaele et al. 2014).

In this study, we aimed to understand the specific effect of the fusion of MBTD1 to EZHIP and delineate whether the molecular mechanism is similar to that of other LGESS fusion proteins, in which NuA4/TIP60 association/activity may prove to be efficient therapeutic targets (Sudarshan et al. 2022; Tavares et al. 2022). Endometrial stromal sarcomas are very rare cancers; therefore, model systems are lacking to study this disease. Moreover, the cell-of-origin of this cancer is poorly described. Accordingly, only heterologous cellular models of LGESS have been developed. The basal chromatin context of the cell-of-origin is consequential to the effects of a disrupted chromatin modifier. Therefore, we performed histone modification CUT&RUN assays in LGESS patient samples with JAZF1-SUZ12 chromosomal translocation. We aimed to understand whether the molecular mechanisms of the JAZF1-SUZ12 fusion described in heterologous cell systems are conserved in patient samples and find other dependencies.

## RESULTS

### Characterization of the Interactome of MBTD1-EZHIP Fusion Protein

MBTD1-EZHIP chromosomal translocation is recurrently found in LGESS, producing a fusion protein that retains amino acids 1 to 589 of MBTD1, including the Zinc Finger and MBT domains I-IV (Dewaele et al. 2014). MBTD1 interacts with the NuA4/TIP60 complex through MBT-I and MBT-II domains and binds to H4K20me1/2 (Jacquet et al. 2016). The fusion retains the EZHIP amino acids 254 to 503, which contains the 12 amino acid KLP domain (K27 M-like peptide) that mediates binding to PRC2 and inhibits EZH2 activity (Jain et al. 2019) (**Figure 1A**). We created isogenic cell lines in K562 cells expressing MBTD1-EZHIP or EZHIP by inserting the gene of interest with a C-terminal 3XFLAG and 2XStrep tag at the AAVS1 safe harbor locus (Dalvai et al. 2015). We then performed tandem affinity purification from selected clones to get homogenous-purified complexes. Mass spectrometry and western blotting were used to determine the identity of the proteins in the purified complexes (**Figure 1B, C**). The MBTD1-EZHIP fusion protein can assemble all NuA4/TIP60 complex components expressed in K562 cells. The fusion protein (as well as EZHIP) interacts with all the core components of the PRC2 complex (**Figure 1B, C**). Interestingly, we found that MBTD1-EZHIP and EZHIP preferentially associate with subunits specific to the PRC2.1 subcomplex, such as PHF19, MTF2, and PHF1 (**Figure 1C**), with no detected peptides for PRC2.2 specific JARID2 and AEBP2 (**Figure 1B**). This preference is not due to the difference in the expression of PRC2 subunits in K562 since the purification of endogenously tagged EZH2 in this cell line showed interaction with both PRC2.1 and 2.2 subunits (published in (Dalvai et al. 2015)) and used for western blotting in **Figure 1B**). The fusion of the NuA4/TIP60 complex to the PRC2.1 complex by MBTD1-EZHIP is reminiscent of the chimeric complexes formed by the other LGESS fusion proteins, EPC1-PHF1 and JAZF1-SUZ12 (Sudarshan et al. 2022). We tested whether MBTD1-EZHIP is a transcriptional activator similar to EPC1-PHF1 and JAZF1-SUZ12 using a dCAS9-based induced recruitment assay (Alerasool et al. 2022). Indeed, MBTD1-EZHIP drives the expression of the GFP reporter to a level comparable to that of the N-terminal portion of MBTD1 present in the fusion (**Figure 1D**). Thus, the MBTD1-EZHIP fusion protein can assemble a chimeric complex of NuA4/TIP60 and PRC2.1 and act as a transcriptional activator. We then took advantage of published separation-of-function mutants of MBTD1 and EZHIP (Jani et al. 2019) and (Zhang et al. 2020), respectively) and created mutations in either the MBTD1 or EZHIP portions of the fusion protein in such a way that it could assemble either NuA4/TIP60 or PRC2.1 (**Figure 1E**). These mutations allowed us to distinguish the effects of these complexes. The A236D, L259D, and L263D mutations in MBTD1 (labelled 3mut henceforth) (Zhang et al. 2020) completely abolished the interaction between the MBTD1-EZHIP fusion protein and KAT5 (NuA4/TIP60 complex) (**Figure 1E, F**). The M406E mutation of EZHIP significantly reduced the interaction between MBTD1-EZHIP and EZH2 (PRC2 complex) (**Figure 1E, F**). The M406E mutation of the fusion protein shows residual interaction with EZH2, suggesting additional interaction mechanisms at play between EZHIP and PRC2. Nevertheless, the M406E mutation disrupted the inhibitory effect of EZHIP on EZH2 (Jani et al. 2019).

**Figure 1.**
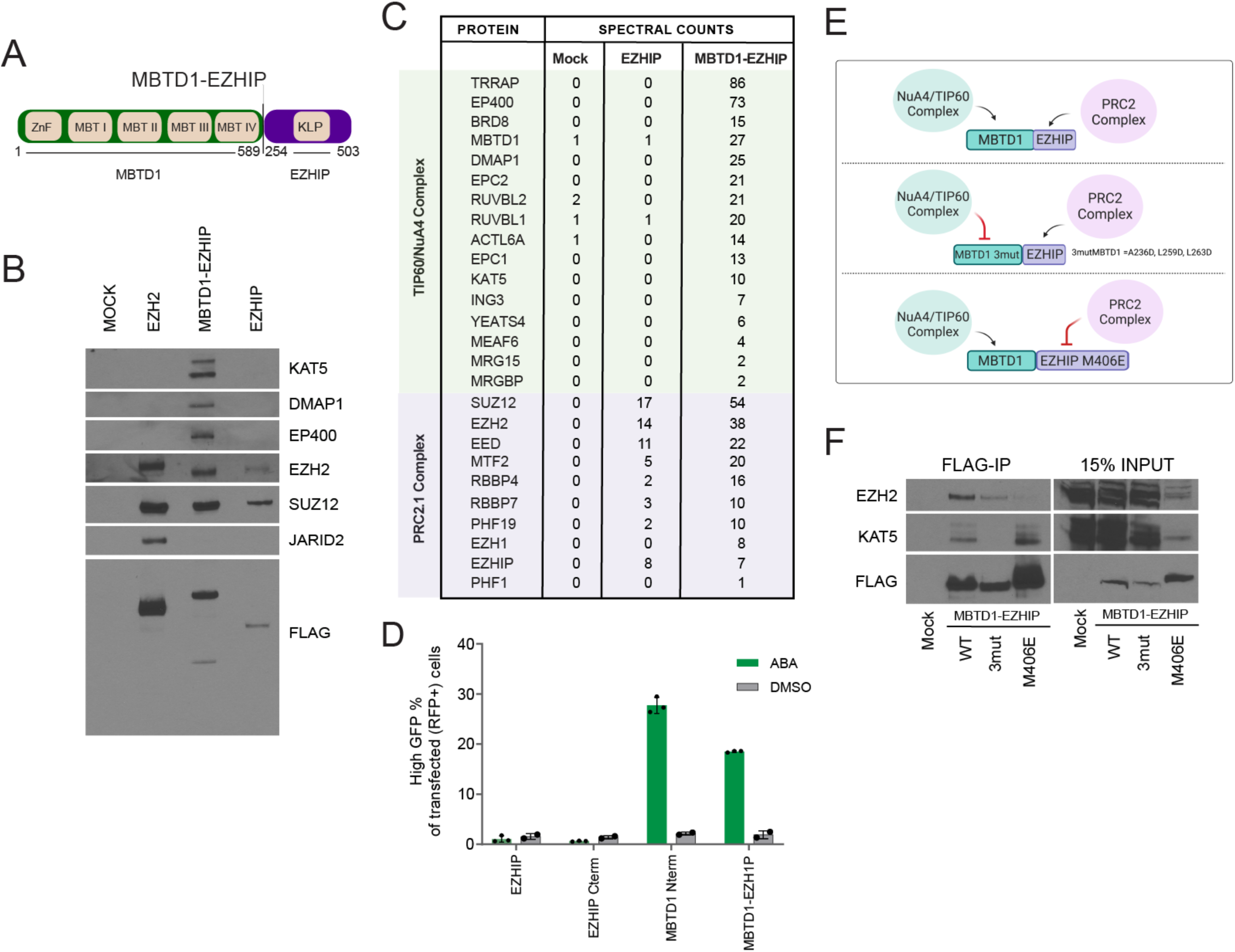
Characterization of the Interactome of the MBTD1-EZHIP Fusion Protein. **(A)** Schematic representation of the MBTD1-EZHIP fusion protein. The numbers indicated are amino acids, protein domains retained in the fusion are indicated**. (B)** Western blots of selected NuA4/TIP60 and PRC2 complex subunits on the affinity purified fractions of the proteins indicated. EZH2 purification is from (Dalvai et al. 2015). **(C)** Mass Spectrometry analysis of the affinity purified complexes shown in (B). **(D)** dCas9-based inducible reporter gene activation assay described in (Alerasool et al. 2022), transcription activation was quantified by Flow cytometry analysis, the % of cells transfected with indicated genes expressing high GFP upon ABA treatment is plotted. At least 25,000 cells were analyzed for each replicate. Error bars represent SEM of 5 independent repeats. DMSO is used as a negative control. **(E)** Schematic representation of the chimeric complex assembled by the MBTD1-EZHIP fusion protein and the mutations that can inhibit such association (Zhang et al. 2020; Jain et al. 2019). **(F)** Western blotting of Flag immunoprecipitation of MBTD1-EZHIP fusion protein and the indicated mutants.

### The MBTD1-EZHIP Fusion Complex Binds at Polycomb Target Genes and Induces Global Changes in Histone Modifications

LGESS fusions were localized to NuA4/TIP60 and PRC2-regulated chromatin regions (Sudarshan et al. 2022). We generated K562 cell lines expressing MBTD1-EZHIP or mutants with a 3XHA tag to localize the MBTD1-EZHIP fusion protein by CUT&RUN profiling. We developed a K562 cell line with an endogenous 3XHA tag at the C-terminus of the EPC1 alleles using CRISPR-cas9. This cell line was used to identify the occupancy of NuA4/TIP60 (anti-EPC1-HA CUT&RUN) and PRC2 (anti-SUZ12 CUT&RUN). The anti-HA tag CUT&RUN revealed the chromatin occupancy of MBTD1-EZHIP and its mutants **(Figure 2A, E, F, G**). The MBTD1-EZHIP fusion complex was found mainly in the intergenic (40.3%) and intronic regions (29.2%) **(Figure 2E),** reflecting the binding pattern of Polycomb complexes **(Figure S1J)**. The M406E mutant primarily bound to the proximal promoter (24.8%), 5’-UTR (17.8%), and introns (17.7%) **(Figure 2F),** reflecting the binding pattern of NuA4/TIP60 in active chromatin regions **(Figure S2K)**. 3mut was expressed at low levels **(Figure 2D)** and showed few binding sites (**Figure 2A, Figure S2C**), mainly in the proximal promoter (33.5%) and 5’-UTR (15.8%) (**Figure 2G**). MBTD1-EZHIP peaks overlapped with PRC2 sites, that is, SUZ12 CUT&RUN peaks in K562 (21.3%), whereas the M406E mutant showed overlap with NuA4/TIP60 sites, that is, EPC1-HA CUT&RUN peaks (49.13%) (**Figure 2H, I**). Thus, the MBTD1-EZHIP fusion complex binds mainly to PRC2-regulated regions and some NuA4/TIP60 regions. Conversely, the M406E mutant showed a loss of binding to the PRC2 regions and gained binding at the NuA4/TIP60 regions. Overall, the anti-HA CUT&RUN data suggest that the fusion of MBTD1 with EZHIP redistributes its binding to the PRC2-bound areas.

**Figure 2.**
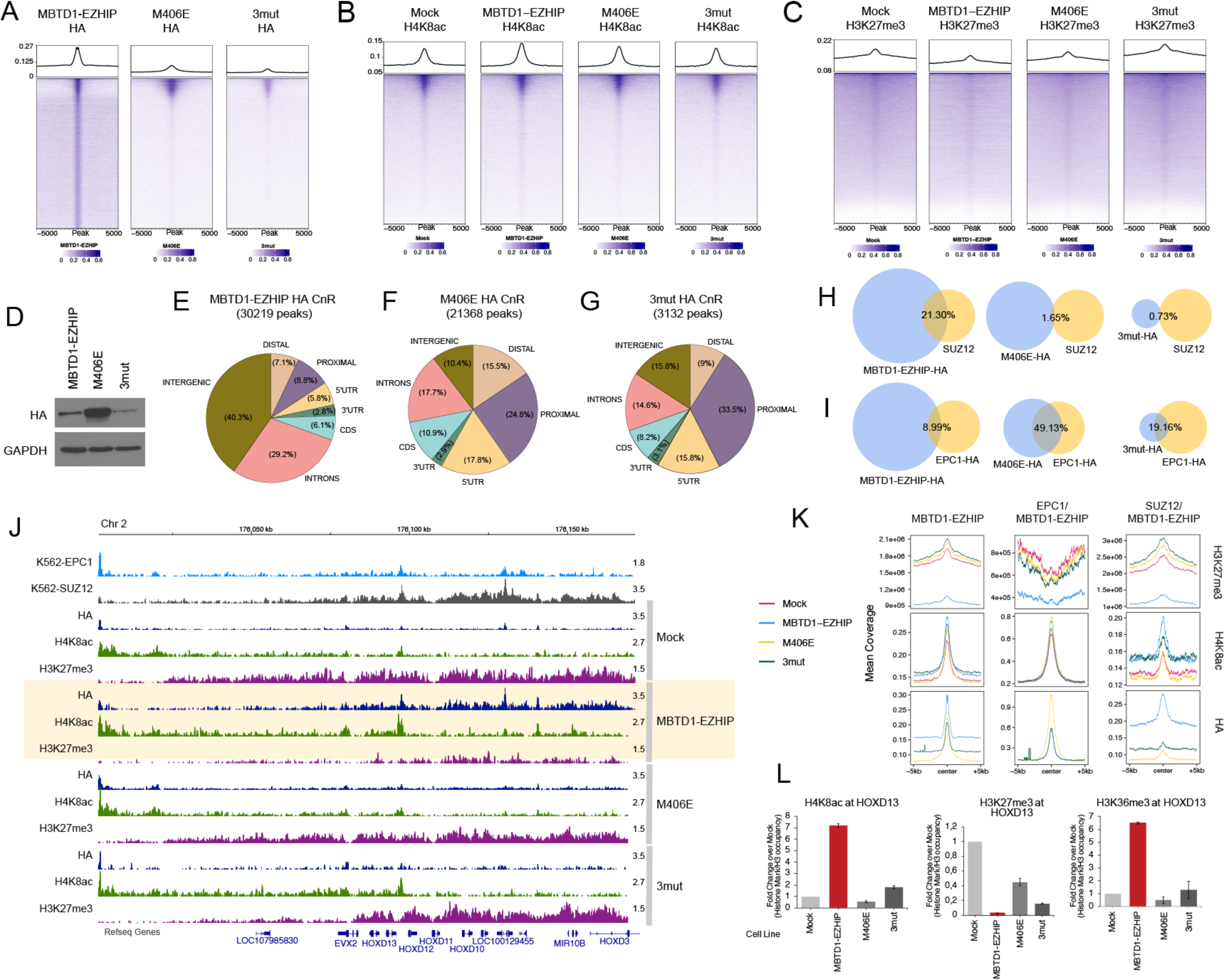
The MBTD1-EZHIP Fusion Complex Induces Global Changes in Histone Modifications. **(A)** Anti-HA CUT&RUN signal in the indicated K562 cell lines. The heat maps show coverage of anti-HA CUT&RUN signal of MBTD1-EZHIP, M406E and 3mut cell lines after subtracting the background from Mock-HA (Negative control), plotted on peaks of MBTD1-EZHIP (± 5000 bp to peak center). **(B)** Anti-H4K8ac CUT&RUN signal in the indicated K562 cell lines. The heat maps show coverage of anti-H4K8ac CUT&RUN signal in Mock (Negative control), MBTD1-EZHIP, M406E and 3mut cell lines plotted on peaks of MBTD1-EZHIP (± 5000 bp to peak center). **(C)** Anti-H3K27me3 CUT&RUN signal in the indicated K562 cell lines. The heat maps show coverage of anti-H3K27me3 CUT&RUN signal in Mock (Negative control), MBTD1-EZHIP, M406E and 3mut cell lines plotted on peaks of MBTD1-EZHIP (± 5000 bp to peak center). **(D)** Protein expression level of MBTD1-EZHIP and mutant (M406E and 3mut) HA tagged K562 cell line clones used in this study. **(E)** MBTD1-EZHIP-HA (F) M406E-HA and (G) 3mut-HA CUT&RUN peaks overlapping with functional elements in the genome. **(H)** Venn diagram showing the overlap of MBTD1-EZHIP-HA, M406E-HA and 3mut-HA CUT&RUN peaks with SUZ12 CUT&RUN peaks in K562. **(I)** Venn diagram showing the overlap of MBTD1-EZHIP-HA, M406E-HA and 3mut-HA CUT&RUN peaks with EPC1-HA CUT&RUN peaks in K562. **(J)** Representative CUT&RUN coverage tracks of HA, H4K8ac and H3K27me3 in Mock, MBTD1-EZHIP, M406E and 3mut K562 cell lines at the HOXD gene cluster. EPC1-HA and SUZ12 CUT&RUN in K562 cells are also shown. Highlighted region in yellow shows enriched HA signal, H4K8ac increase and reduction in H3K27me3 in MBTD1-EZHIP-HA expressing K562 cell line. **(K)** Aggregate density plots (Mean coverage) of HA, H4K8ac and H3K27me3 CUT&RUN in indicated cell lines at peaks that are common between MBTD1-EZHIP-HA and SUZ12 as well as MBTD1-EZHIP-HA and EPC1-HA. **(L)** H4K8ac, H3K27me3 and H3K36me3 ChIP-qPCR in Mock, MBTD1-EZHIP, M406E and 3mut K562 cell lines at the HOXD13 gene. Bar graph in maroon color highlights the results in the MBTD1-EZHIP cell line. (Values are a ratio of %Input of indicated histone modification and H3; Graphs show the fold change over Mock/negative control) (n=2, error bars are range of the values).

The *HOXD* gene cluster in K562 cells showed binding of SUZ12 (grey peaks) and EPC1 (blue peaks) (**Figure 2J**). At this locus, we observed the binding of MBTD1-EZHIP (navy blue peaks), mirroring the binding of SUZ12 (MBTD1-EZHIP cell line highlighted in yellow) (**Figure 2J**). The regions bound by MBTD1-EZHIP also showed a drastic reduction in H3K27me3 levels (purple peaks) and an increase in H4K8ac levels (green peaks) (see also aggregate density plots at MBTD1-EZHIP peaks in **Figure 2K**). ChIP-qPCR of the *HOXD13* gene confirmed these observations and showed increased transcription-linked H3K36me3 histone modification (**Figure 2L**).

### The MBTD1-EZHIP Fusion Complex is Dependent on both NuA4/TIP60 and PRC2 Complexes to Upregulate Gene Expression

Changes in histone modifications, such as H4K8ac and H3K27me3, induced by the MBTD1-EZHIP fusion complex are indicators of underlying changes in gene expression. We overexpressed MBTD1-EZHIP in K562 cells using a lentiviral system. MBTD1-EZHIP expressing cells compared to non-expressing cells showed predominantly upregulated genes (**Figure 3A**). We selected a set of these upregulated genes (*HOXD13, NOTCH2, BMP6, RPSA* as control) and tested their expression by RT-qPCR in K562 cell lines expressing the MBTD1-EZHIP fusion protein or mutants from the AAVS1 locus **(Figure 3B, C**). M406E and 3mut cell lines did not show upregulation in the MBTD1-EZHIP cell line. Thus, NuA4/TIP60 and PRC2 are important for gene expression changes induced by the fusion oncoprotein. Furthermore, although we observed minor changes in histone modifications in M406E and 3mut cell lines at the *HOXD13* gene (**Figure 2J, L**), it did not result in the upregulation of gene expression, demonstrating the synergistic effect of H3K27me3 loss and H4/H2A acetylation gain. We observed an enrichment of the coverage of MBTD1-EZHIP-HA CUT&RUN on upregulated genes when compared to mutants or controls (**Figure 3D**), indicating that MBTD1-EZHIP directly regulates these genes. Moreover, the upregulated genes also showed enrichment for SUZ12 and modest binding of EPC1-HA, further supporting the hypothesis that MBTD1-EZHIP binds to Polycomb target genes with some bivalency and upregulates them (**Figure 3D**).

**Figure 3.**
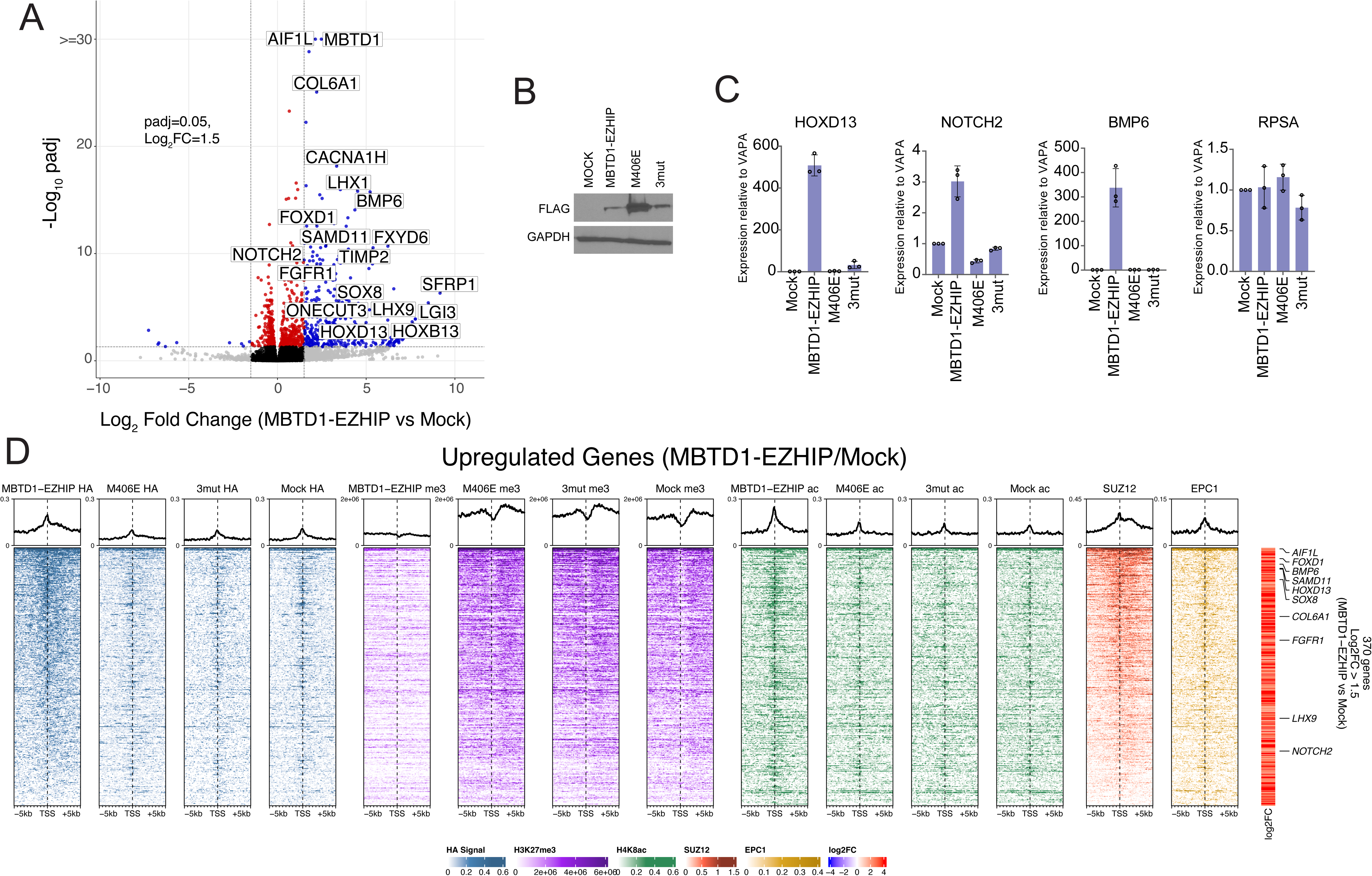
MBTD1-EZHIP Fusion Complex is Dependent on both NuA4/TIP60 as well as PRC2 Complexes to Upregulate Gene Expression. **(A)** RNA-sequencing followed by differential expression analysis, represented here as a Volcano plot. K562 cells expressing MBTD1-EZHIP fusion is compared to empty vector K562 cells. The graph shows a cut off at 0.05 for padj value and at 1.5 for Log2 fold change. An upper limit for - Log10 padj value was set at 30 at the Y axis. **(B)** Western blotting showing the expression of the fusion protein and mutants in cell lines used in (C). **(C)** RT-qPCR at selected upregulated genes (*HOXD13, BMP6, NOTCH2*) from (A), performed in K562 cell lines expressing MBTD1-EZHIP or mutants compared to an empty vector control. Expression level is plotted relative to the gene VAPA. The RPSA gene is a constantly expressed gene used as control. (n=3, error bars are standard deviation). **(D)** Integration of CUT&RUN and RNA sequencing: Heatmap and aggregate plots of anti-HA, SUZ12, H3K27me3 and H4K8ac CUT&RUN plotted at upregulated genes (± 5000 bp from TSS) from (A).

The MBTD1-EZHIP-mediated decrease in H3K27me3 and increase in H4K8ac occupancy observed genome-wide (**Figure 2B, C**) was striking when plotted on upregulated genes in MBTD1-EZHIP (**Figure 3D**). Neither M406E nor 3mut showed occupancy or changes in histone modifications in genes upregulated by MBTD1-EZHIP (**Figure 3D**), revealing that MBTD1 (NuA4/TIP60) is not required for MBTD1-EZHIP occupancy. Unfortunately, the low expression of 3mut obscured its binding to the SUZ12/PRC2 and MBTD1-EZHIP co-occupied regions. Regardless, our data with the MBTD1-EZHIP CUT&RUN show that PRC2 influences its genome occupancy and that the decrease in H3K27me3 and gain of H4K8ac (through NuA4/TIP60) are the predominant mechanisms employed by the fusion of upregulated genes.

### Conserved Gene Expression Pattern between JAZF1-SUZ12 and MBTD1-EZHIP

Previous studies have suggested clustering of gene expression patterns and similarities in the histological manifestation of the disease between JAZF1-SUZ12 and MBTD1-EZHIP translocated LGESS (Dewaele et al. 2014). We further explored this phenomenon by overexpressing either MBTD1-EZHIP or JAZF1-SUZ12 in cells that may resemble the cell-of-origin of LGESS, that is, mesenchymal stem cells-bmMSC (bone marrow-derived and immortalized by hTERT) and human immortalized endometrial stromal cells-HIESC (immortalized by SV40 Large T-antigen). We performed RNA sequencing on these cells to understand their similarities and differences—both MBTD1-EZHIP and JAZF1-SUZ12 mediated differential gene expression when compared to the control. Similar to experiments in K562 cells (Sudarshan et al. 2022) and **Figure 3A**), gene upregulation was predominant in fusion-expressing bmMSCs and HIESCs. A similar set of genes was upregulated in JAZF1-SUZ12 and MBTD1-EZHIP expressing bmMSC (**Figures 4A-C & S2A-B**) or HIESC cells (**Figure S3A-C, J**). Over-representation analysis predicted that PRC2 components such as EZH2 and SUZ12 (**Figures 4D and S3H-I**) and PRC2-mediated H3K27me3 histone modification (**Figures 4E and S3F-G**) bind the upregulated genes. This prediction is consistent with the oncogenic mechanisms of JAZF1-SUZ12 (Sudarshan et al. 2022; Tavares et al. 2022) and MBTD1-EZHIP (**Figures 1–3**), being the upregulation of Polycomb target genes.

**Figure 4.**
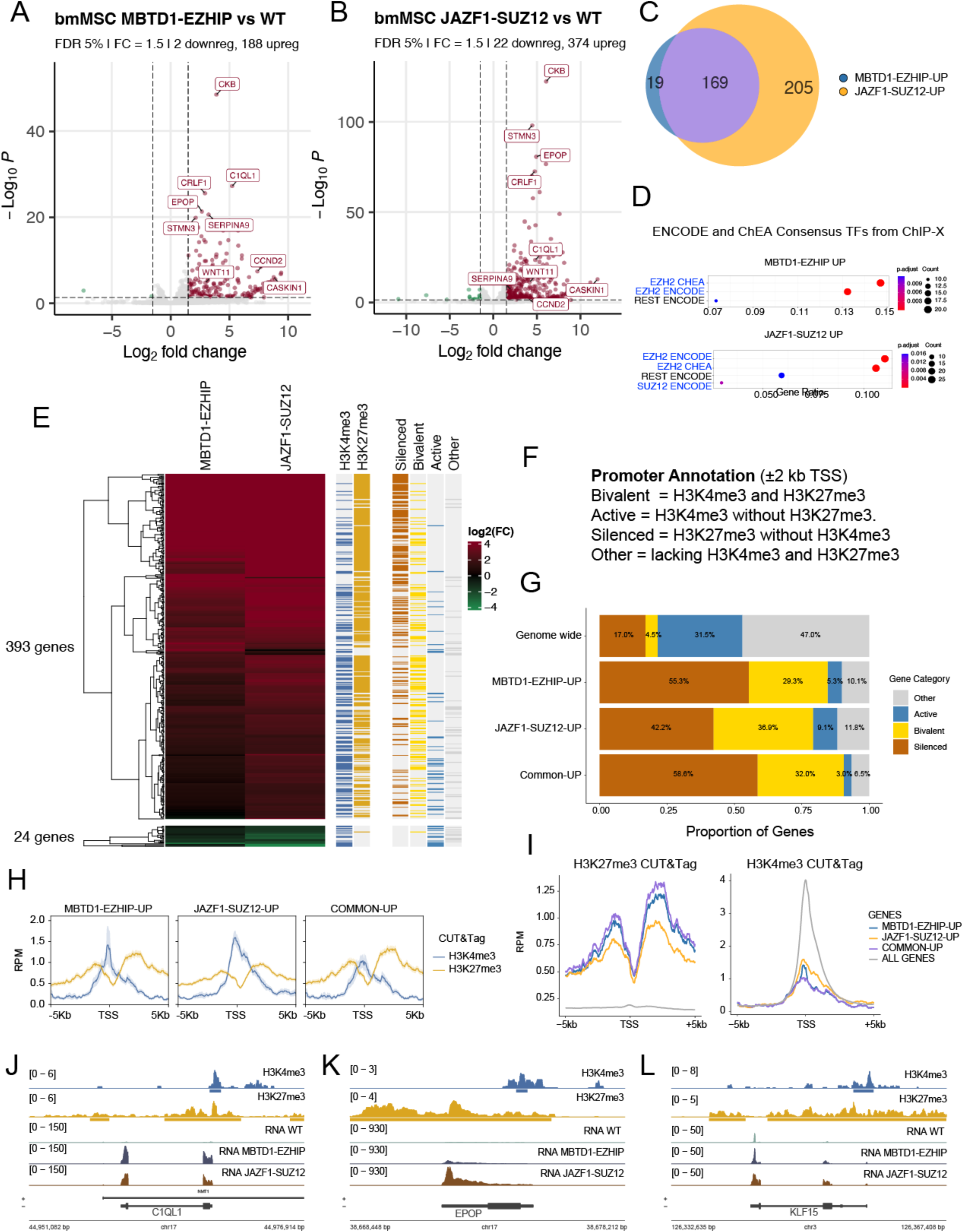
Conserved Gene Expression Patterns between JAZF1-SUZ12 and MBTD1-EZHIP. **(A)** RNA-sequencing Volcano plot of bmMSC cells expressing MBTD1-EZHIP fusion protein compared to empty vector control/Mock. The graphs show a cut-off at 0.05 for the padj value, and 1.5 for Log2 fold change. An upper limit for -Log10 padj value was set at 80 and 30 at the Y axis, respectively. **(B)** Volcano plot of bmMSC cells expressing JAZF1-SUZ12 fusion protein compared to empty vector control/Mock. **(C)** Venn diagram showing the significant overlap between upregulated genes in JAZF1-SUZ12 and MBTD1-EZHIP expressing bmMSC cells (Log2FC 1.5). **(D)** Gene over-representation analysis of upregulated genes using the ENCODE_and_ChEA_Consensus_TFs_from_ChIP-X. **(E)** Heatmap correlating differentially regulated genes by MBTD1-EZHIP and JAZF1-SUZ12 fusions in bmMSC and CUT&TAG mapping of H3K4me3 and H3K27me3 in these cells. **(F-G)** Promoter annotations as bivalent, active, silenced and others based on presence of H3K4me3 and/or H3K27me3, and analysis of the ones up-regulated by the fusions compared to genome-wide promoters. **(H-I)** Aggregate plots of H3K4me3 and H3K27me3 CUT&TAG plotted at promoters of genes upregulated by the fusions. **(J-L)** Representative CUT&TAG tracks of H3K4me3 and H3K27me3 in bmMSC cells at genes upregulated by the MBTD1-EZHIP and JAZF1-SUZ12 (*C1QL1*, *EPOP* and *KLF15*). RNA-seq tracks are also shown.

GSEA analysis with the Hallmark pathway gene set (M.Sig.Db) in upregulated genes in bmMSCs (expressing JAZF1-SUZ12 or MBTD1-EZHIP) showed enrichment of genes downregulated by KRAS activation (KRAS signaling Down) as well as genes involved in myogenesis (**Figure S2D-E**). Upregulation of the myogenesis pathway is interesting because some LGESS tumors indeed show muscle differentiation (Huang, Ladanyi, and Soslow 2004). In HIESC, pathways that showed enrichment in upregulated genes were WNT signaling pathway genes, genes upregulated during angiogenesis, genes downregulated by KRAS activation, and genes defining late response to estrogen (**Figure S3D-E**). Gene Ontology terms (GO-Biological processes) such as neuronal differentiation, were enriched in both bmMSC and HIESC cells expressing JAZF1-SUZ12 or MBTD1-EZHIP (**Figures S2F-G and S3K-L**). Importantly, JAZF1-SUZ12 or MBTD1-EZHIP-mediated differential gene expression in HIESC matched the previously reported gene expression changes in primary endometrial stromal cells with JAZF1-SUZ12 expression (Tavares et al. 2022). Thus, MBTD1-EZHIP converged the perturbed oncogenic gene expression pattern with JAZF1-SUZ12, suggesting that a difference in the finer molecular mechanism may still produce similar phenotypic outcomes in LGESS with various fusion proteins. CUT&TAG analysis of bmMSC cells for H3K4me3 and H3K27me3 marks allowed us to annotate gene promoters as active (H3K4me3 alone), silent (H3K27me3 alone) or bivalent (both). Analysis of genes upregulated by JAZF1-SUZ12 and MBTD1-EZHIP shows not only a strong enrichment for silent promoters but even a stronger one for bivalent promoters (**Figures 4E-G, S2C**). This link to bivalent promoters being targeted by the fusions for activation is well demonstrated by metaplots as well as genome browser views of normalized read counts on specific genes (CUT&TAG and RNA-seq) (**Figure 4H-L**).

### JAZF1-SUZ12 Expressing Patient Tumors Show Redistribution of Histone Modifications

We previously reported the transcriptomics of LGESS patient samples with JAZF1-SUZ12 fusion and a paired normal endometrial sample. Pairwise differential expression analysis of these samples revealed significantly upregulated and downregulated genes (Sudarshan et al. 2022)). To understand the histone modification changes mediated by JAZF1-SUZ12 (through NuA4/TIP60 and PRC2) and their correlation with the observed gene expression changes, we performed H4K8ac and H3K27me3 CUT&RUN profiling on these patient samples. **Figure 5A** shows the samples used in the (Sudarshan et al. 2022) and this study. We plotted the coverage of histone modifications at differentially expressed genes in JAZF1-SUZ12 translocated endometrial tumor compared to matched normal endometrial tissue. We observed an increase in H4K8ac occupancy and a decrease in H3K27me3 at upregulated genes in endometrial and uterine tumor samples compared to those in normal tissue (heatmaps/metaplots/boxplots, **Figure 5B-D**). We also presented the coverage of H3K27me3 and H4K8ac at selected upregulated genes, such as the *HOXA* gene cluster (*HOXA9, HOXA10*, and *HOXA11*) (**Figure 5E-F**), *HOXD* gene cluster (*HOXD9, HOXD10*, and *HOXD11*) (**Figure 5E, H**), *WNT6* (**Figure S4A, D**), and *SAMD11* (**Figure S4A, E**).

**Figure 5.**
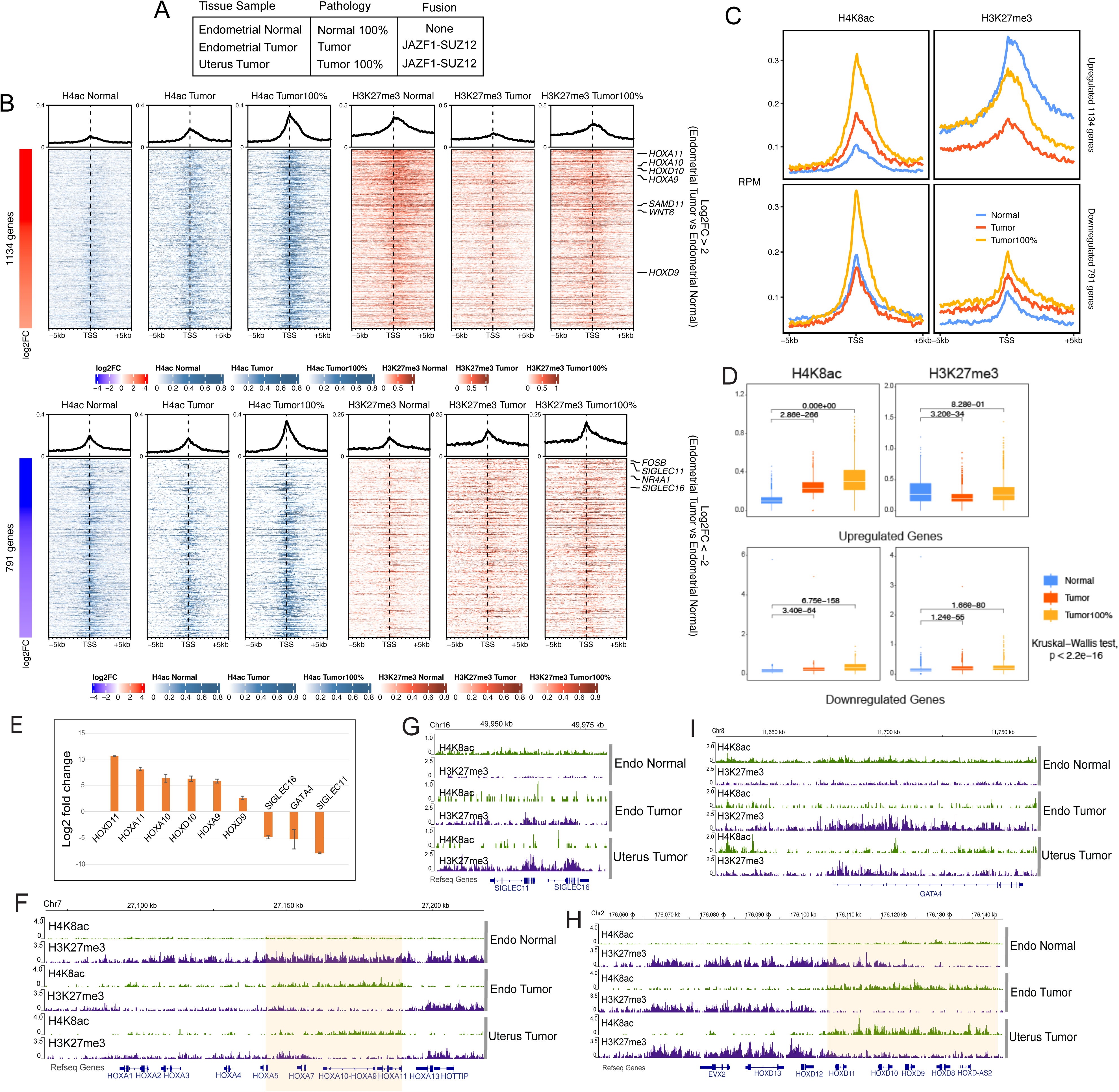
JAZF1-SUZ12 expressing patient tumors show redistribution of histone modifications. **(A)** Details of the patient samples used in the study (see also Sudarshan et al. 2022). **(B-D)** Heatmap, aggregate and box plots of anti-H4K8ac and H3K27me3 CUT&RUN signal in the indicated patient sample. The CUT&RUN coverage is plotted on upregulated or downregulated genes (± 5000 bp from TSS) in endometrial tumor compared to normal tissue. **(E)** Selected genes showing change in expression level in JAZF1-SUZ12 expressing patient tumors reported in (Sudarshan et al. 2022). **(F-I)** Representative anti-H4K8ac and anti-H3K27me3 CUT&RUN coverage tracks at differentially expressed genes (shown in B) in normal and patient tumor tissue samples.

Surprisingly, we observed an increase in H3K27me3 at downregulated genes in endometrial and uterine tumor samples compared to normal tissues (**Figure 5B-D**). Coverage of H3K27me3 and H4K8ac at selected downregulated genes, such as *SIGLEC11*, *SIGLEC16* (**Figure 5E, G**), *GATA4* (**Figure 5I**), *NR4A1* (**Figure S4A-B**), and *FOSB* (**Figure S4A, C**) are also shown. The average coverage of H4K8ac in downregulated genes was inconsistent between endometrial and uterine tumors (**Figure 5B-D)**. Furthermore, the IGV tracks at selected genes (**Figures 5G, I, and S4B-C**) showed low CUT&RUN signals, which precludes us from concluding H4K8ac occupancy levels at downregulated genes. However, no significant decrease was observed.

There are two possibilities for the increase in H3K27me3 at downregulated genes: (1) direct binding of the fusion protein at active genes and deposition of H3K27me3, and (2) indirect redistribution of H3K27me3 loss at the fusion protein-bound genes. Although normalized genome-wide correlation of gene expression and histone modifications in our LGESS samples is needed to reveal more details, it is interesting to note that gene over-representation analysis of both upregulated and downregulated genes predicted that these genes are regulated by H3K27me3 (Sudarshan et al. 2022). This evidence supports the model of indirect redistribution of H3K27me3 to regions that are active/de-repressed Polycomb targets. Moreover, redistribution of H3K27me3 and silencing of tumor suppressors is a common mechanism in cancers that perturb the genome-wide levels of H3K27me3 such as H3K27M mutated HGG and EZHIP-overexpressed PFA Ependymoma (Bender et al. 2013; Harutyunyan et al. 2019; Stafford et al. 2018; Larson et al. 2019; Mohammad et al. 2017; Jain et al. 2019; Chan et al. 2013). Unfortunately, we lack a JAZF1-SUZ12 fusion protein-specific antibody to map its chromatin localization, which could have given us definitive proof of redistribution of H3K27me3.

## DISCUSSION

Our study confirmed not only the interactome of the MBTD1-EZHIP fusion protein but demonstrates clearly it leads to the formation of a stable hybrid NuA4/TIP60-PRC2 complex. Our data also revealed its effect on histone modifications and gene expression in cells. Although EZHIP has been reported to interact with EZH2 and purify PRC2.1 and PRC2.2 (Jain et al. 2019), our study with EZHIP and MBTD1-EZHIP showed a preference for PRC2.1. This discrepancy could be attributed to the difference in protein expression level observed in different studies. Since our experiments utilized the insertion of the gene of interest at the AAVS1 safe harbor locus and used a PGK1 promoter, protein expression was maintained at lower levels. This system has been previously used to purify stoichiometric complexes and thus may provide better clarity on the composition of the complexes (Dalvai et al. 2015).

Furthermore, another study reported the preference of EZHIP for PRC2.1 (Ragazzini et al. 2019). Interestingly, recent data shows that PRC2.1 is more important for maintaining bivalent domains in mESCs than the PRC2.2 sub-complex (Perino et al. 2020) and a recent study showed that PRC2.1 is responsible for depositing the bulk of H3K27me3 in cells (Glancy et al. 2023); thus, studying the mechanisms for the preference of EZHIP for PRC2.1 could yield important insights into the effect of PRC2 inhibitory proteins.

In this study, we utilized mutants to clarify the contribution of EZHIP/PRC2.1 and NuA4/TIP60 to the oncogenic effect of MBTD1-EZHIP expression. We inferred the impact of the loss of EZHIP activity in MBTD1-EZHIP through the M406E mutant, which behaves like MBTD1 and occupies regions bound by NuA4/TIP60. The M406E mutant increased H4K8ac at bound regions but showed no decrease in H3K27me3 and, significantly, did not bind and upregulate MBTD1-EZHIP target genes. Thus, EZHIP is critical for the localization of the fusion protein, and its function in inhibiting EZH2 is essential for gene activation.

Our epigenomic data in JAZF1-SUZ12 translocated LGESS patient samples showed, for the first time, an increase in H3K27me3 in downregulated genes compared to normal endometrial tissue. Thus, in addition to upregulating oncogenes, the increase in H3K27me3 levels implicates the JAZF1-SUZ12 fusion protein in suppressing genes that may play an essential role in LGESS tumorigenesis. Interestingly, the top downregulated genes in our LGESS patient samples were important immune response genes that are involved in immune clearance during menstrual shedding This is in agreement with an earlier study modeling JAZF1-SUZ12 during the decidualization of primary endometrial stromal cells (Tavares et al. 2022; B. Xu et al. 2014). Thus, the fusion proteins in LGESS may maintain the endometrial stromal cells in a proliferative phase by disrupting various aspects of the decidualization/differentiation process via mislocalization of histone marks, leading to tumorigenesis (Tavares et al. 2022).

Based on these findings, EZH2 and Tip60/KAT5 inhibitors may be useful as potential therapeutic approaches. Alternatively, based on our results with MBTD1-EZHIP mutants, small-molecule inhibitors that can disrupt the interaction of the fusion protein with NuA4/TIP60 and PRC2 complexes can be viable therapeutic options.

### Limitations

Low expression of the mutant separating MBTD1-EZHIP from the NuA4/TIP60 complex (3mut) prevents us from fully understanding the role of NuA4/TIP60 interaction with MBTD1-EZHIP. This mutant would also have provided us with information about the difference in the misexpression of EZHIP compared to the formation of the MBTD1-EZHIP fusion protein. Thus, there is a need to study MBTD1 separation mutant (3mut) in comparison to MBTD1-EZHIP in a different cellular background or using an inducible expression system.

LGESS is a rare cancer that lacks a sound model system to understand oncogenic mechanisms fully. Patient samples are a precious and limited resource, which prevented further exploration of the mechanism of H3K27me3 upregulation in LGESS. We predict that the decidualization of endometrial stromal cells expressing TIP60-PRC2 fusions would provide the right chromatin background to study LGESS. *JAZF1* and *MBTD1* also play known roles in endometrial decidualization (Chadchan et al. 2020; Liang et al. 2023). Therefore, the presence of the endogenous intact allele of *JAZF1* or *MBTD1* in cell line models may have obscured subtle histone modifications and gene expression changes in our study. Hence, creating TIP60-PRC2 chromosomal translocations using CRISPR-cas9 technology in endometrial stromal cells may provide a better model system for understanding LGESS.

## Materials and Methods

### Cell culture

K562 cells were obtained from the ATCC and maintained at 37°C under 5% CO2 in RPMI medium supplemented with 10% fetal bovine serum/newborn calf serum and GlutaMAX. HEPES-NaOH (25 mM, pH 7.4) was added during the culture in the Spinner flasks. HIESC cells immortalized with SV40 large T antigen were a kind gift from Dr. Michel A. Fortier and Yannick Doyon. Cells were cultured as previously described (Chapdelaine et al. 2006). Bone marrow-derived Mesenchymal Stromal cells (bmMSCs) immortalized with *hTERT* were a kind gift from Dr. Nada Jabado. The cells were maintained at 37°C under 5% CO2 in low-glucose DMEM (Thermo Scientific 10567022) supplemented with 5% Stemulate (Sexton Biotechnologies).

### Patient tissue samples

The Research Ethics Board of the University Health Network in Toronto, ON, Canada, approved this study. Biobanked low-grade endometrial stromal sarcoma specimens (frozen) were obtained with broad consent. Fluorescence in situ hybridization (FISH) was performed to identify rearrangement involving JAZF1 and SUZ12.

### Construction of recombinant DNA

Prof. Nada Jabado provided the EZHIP ORF construct. The MBTD1 ORF construct and the 3mut version of MBTD1 were obtained from a previous study in the lab (Jani et al. 2019). Primer extension mutagenesis was employed to generate the MBTD1-EZHIP fusion gene and introduce mutations in the MBTD1-EZHIP construct. This involved performing independent nested PCRs using overlapping primers with the desired mutation or chimeric DNA. The resulting PCR products were combined for the final PCR step. The MBTD1-EZHIP, M406E, and 3mut constructs were cloned into the pDONOR223 Gateway cloning plasmid using Gateway BP clonase (Thermo Fisher Scientific). Subsequently, the constructs were transferred to a lentiviral destination vector (Addgene 39481) or custom destination expression vectors using Gateway LR clonase (Thermo Fisher Scientific). The custom destination vectors were designed with AAVS1 homology arms and featured C-terminal tandem 3XFLAG and 2XStrep tags or 3XHA and 2XStrep tags. Custom vectors were modified from Addgene 68375 (Dalvai et al. 2015). All constructs were subjected to Sanger sequencing verification, and their expression was evaluated through transient transfection experiments.

### Cell line generation

MBTD1-EZHIP and its mutants were specifically targeted to the AAVS1 safe harbor locus in K562 cells using zinc finger nucleases, as previously described by (Hockemeyer et al. 2009; Dalvai et al. 2015). For lentiviral overexpression, plasmids containing the genes of interest were co-transfected with vesicular stomatitis virus G (VSV-G) plasmid DNA and Gag-Pol-Tet-Rev plasmid DNA into HEK293T cells, using polyethyleneimine (PEI) as a transfection reagent. After 48–72 h of transfection, the supernatants containing the lentivirus were collected, and target cells were transduced with the lentivirus in the presence of 8 µg/mL hexadimethrine bromide (polybrene; Sigma) at a multiplicity of infection (MOI) of 0.3. The assays were performed 24 hours after transduction.

### Purification of complexes

Native complexes were purified as previously described (Hockemeyer et al. 2009; Doyon and Côté 2016). The purified complexes were loaded on NuPAGE 4–12% Bis-Tris gels (Invitrogen) and visualized by silver staining. The fractions were then analyzed using mass spectrometry.

### Mass Spectrometry Analysis

The purified fractions were loaded on a 10% SDS-PAGE, run until the dye migrated approximately 1 cm, stained with Sypro Ruby Red, and a gel slice containing the entire protein signal was cut and processed for in-gel digestion with trypsin. Mass spectrometry (MS) analyses were performed using the Proteomics Platform of the CHU de Québec-Université Laval Research Center. The samples were analyzed by nano-LC/MS–MS, either with a 5600 triple TOF or an Orbitrap Fusion. Database Searching Mascot generic format peak list files were created using Protein Pilot, version 4.5 software (Sciex) for the data obtained using the 5600+ triple TOF and Proteome Discoverer 2.3 software (Thermo) for the Orbitrap data. The Mascot generic format sample files were then analyzed using Mascot (Matrix Science; version 2.5.1). Mascot was set up to search a contaminant database and UniProtKB Homo sapiens database, assuming the digestion enzyme trypsin. Scaffold (version Scaffold_4.8.7; Proteome Software, Inc.) was used to validate MS/MS-based peptide and protein identification. Data were compared to mock-purified fractions obtained from cells expressing an empty TAP tag at the AAVS1 site. Data were further analyzed using the CRAPome online tool and filtered against similar experiments (www.crapome.org).

### FLAG Immunoprecipitation

K562 cells were cultured and expanded to obtain a cell count ranging from six to eight million cells. Subsequently, the cells were harvested, subjected to two washes with 1X phosphate-buffered saline (PBS), and then lysed (with buffer containing 450 mM NaCl, 10% glycerol, 50 mM Tris-HCl at pH 8, 1% Triton X-100, 2 mM MgCl2, 0.1 mM ZnCl2, 2 mM EDTA, 1 mM DTT, and protease inhibitors) with twice the volume of the cell pellet, for 30 min at 4°C on an end rotor.

To achieve a final salt concentration of 225 mM, an equivalent volume of lysis buffer without any salt was added to the lysate. The lysate was centrifuged to prepare the whole-cell extracts.

The resulting extracts were incubated with FLAG-M2 agarose resin (Sigma-Aldrich) at 4°C for 4 hours. Following incubation, the resin was separated by centrifugation, washed using lysis buffer containing 225mM NaCl, and subsequently eluted using 3XFLAG peptide (Sigma). The eluted fraction was then loaded onto 4–15% gradient gels along with the input samples and subjected to immunoblotting using appropriate antibodies.

### Chromatin Immunoprecipitation (ChIP)

Histone modification ChIPs were performed according to the protocol described by (Lalonde et al. 2013) with the following modification: The final DNA extraction was performed using the QiaQuick DNA extraction columns following the manufacturer’s instructions (Qiagen) instead of Phenol: Chloroform: Isoamyl alcohol extraction.

Quantitative real-time PCRs were performed on a LightCycler 480 (Roche) with SYBR Green I (Roche) to confirm specific enrichment at the defined loci. Error bars represent the standard error based on two independent experiments. The qRT-PCR primers used are listed in (Sudarshan et al. 2022).

### CUT&RUN sequencing

The experimental procedure followed in this study was based on the protocol outlined by (Skene, Henikoff, and Henikoff 2018) with some modifications.

Frozen patient tissue samples were shattered in a mortar and pestle in a sterile autoclave bag under liquid nitrogen. The shattered tissue was weighed to collect similar amounts of tissue across the samples. The tissue samples were then crosslinked with 0.1% formaldehyde in 1X PBS for 10 min. The reaction was quenched with 125mM Glycine. The cross-linked samples were homogenized in a wash buffer to dissociate the fibrous tissue to obtain a cell suspension.

K562 cells (0.5 million were collected and resuspended in 1X phosphate-buffered saline (PBS). The cells were subjected to crosslinking by treatment with 0.1% formaldehyde for 1 min.

Following cross-linking, the cell suspension was washed at room temperature and incubated with 10 μL of concanavalin-A bead slurry, allowing the cells to bind to the beads. The cell and bead suspensions were incubated overnight at 4°C in buffer containing 0.05% digitonin and 0.5 µg of antibody. The beads were then washed with digitonin buffer and resuspended in a buffer containing pAG-MNase (diluted 1:20; CUTANA EpiCypher) and digitonin, followed by an incubation period of 10 min with agitation. The beads were then washed and resuspended in ice-cold digitonin buffer. CaCl2 (1 mM) was added to the bead suspension to release chromatin, which was then incubated for 2h at 4°C. The reaction was terminated by adding stop buffer containing EDTA and EGTA, followed by an incubation period of 10 minutes at 37°C. The supernatant containing released chromatin was separated from the beads and subjected to overnight de-crosslinking at 55°C. DNA purification was performed using the NEB Monarch PCR and DNA purification kit according to the protocol provided to enrich short DNA fragments. The resulting DNA was quantified using a Qubit HS DNA kit.

The antibodies used were anti-HA antibody (Epicypher 13-2010), Epicypher Rabbit IgG 13-0042, H4K8ac Abcam ab45160, H3K27me3 Cell Sig C36B11.

For assays in K562 cells, the H4K8ac reaction was spiked with 1:50 Drosophila S2 cells, as described previously (Sarthy et al. 2020). The H3K27me3 reaction was spiked with the SNAP-CUTANA K-MetStat Panel by Epicypher, following the manufacturer’s instructions.

The NGS library was prepared according to the manufacturer’s instructions using the NEBNext Ultra II DNA kit for low-input ChIP. One-hundred-base-pair paired-end sequencing was performed using the Illumina NovaSeq 6000 system. Raw reads were trimmed using fastp v0.21.0 (S. Chen et al. 2018). Trimmed reads were aligned to the human genome (hg38) using bwa mem v0.7.17 17 (Heng Li 2013) and SAMtools v1.13 (H. Li et al. 2009). Raw signal tracks and normalized tracks (in reads per million [RPM]) were produced from mapped reads using deepTools’ v2.17.0 bamCoverage tool (F. Ramírez et al. 2014) and BEDTools’ genomecov tool (Quinlan and Hall 2010), respectively. Tracks were converted to the bigwig format using bedGraphToBigWig v2.8 (Kent et al. 2010). MACS2 v2.2.1 software (Feng et al. 2012) performed peak calling. Heat maps were generated using ComplexHeatmap v2.6.2 (Gu, Eils, and Schlesner 2016) and EnrichedHeatmap v1.20.0 packages (Gu et al. 2018), as described for ChIP sequencing data. Peak Overlaps and annotations were performed using the SeqCode Toolkit (Blanco, González-Ramírez, and Di Croce 2021). The CUT&RUN coverage tracks were visualized using an IGV web browser. CUT&TAG on bmMSC cells was performed following the kit manufacturer instructions (CUTANA, Epicypher), and counts were simply normalized by RPMs.

### Recruitment activator assay

This assay was performed as previously described (Alerasool et al. 2022). A HEK293T TRE3G-EGFP reporter cell line was generated by introducing ABI-dCas9 and a guide RNA (gRNA) targeting seven tetO repeats in the TRE3G promoter. A clone that exhibited robust EGFP induction upon stimulation by the potent transcriptional activator VPR was selected for subsequent assays. To initiate the assay, 96-well plates were prepared by seeding 3 × 10^4 cells per well one day before transfection. Each well was transfected with 150 ng of the respective construct using polyethyleneimine (PEI) as the transfection reagent. After 24 h of transfection, the cells were induced by treatment with 100 µM abscisic acid. Following a 48-hour induction period, the cells were dissociated and resuspended in a flow buffer using a liquid handling robot. Subsequently, cell suspensions were analyzed using an LSRFortessa flow cytometer (BD Biosciences). Flow cytometry data were processed using FlowJo software, initially gating for positive gRNA (EBFP2) signals, followed by further gating for construct (TagRFP) expression. A minimum of 25,000 cells were analyzed for each experimental replicate.

### mRNA sequencing

RNA extraction from cells was conducted using the Monarch Total RNA Purification Kit (NEB) following the manufacturer’s protocol. Only samples with an RNA Integrity Number (RIN) greater than 7 were used for library preparation, performed using the NEBNext Ultra II directional RNA library kit in conjunction with the NEBNext Poly (A) mRNA dual-index kit. The subsequent sequencing run was executed on an Illumina NovaSeq 6000 system, generating 100-base-pair paired-end reads. To ensure data quality, the sequencing reads were trimmed using fastp v0.20.1 (S. Chen et al. 2018). Raw and trimmed data quality were assessed using FastQC v0.11.7 and MultiQC v1.5 (Ewels et al. 2016). Kallisto v0.46.2 (Bray et al. 2016) was employed for quantification, with the human genome (hg38) as the reference.

Differential expression analysis was conducted using the DESeq2 v1.30.1 package (Love, Huber, and Anders 2014). Volcano plots representing differential gene expression were generated using the Bioconductor package Enhanced Volcano. Gene Set Enrichment Analysis (GSEA) and overrepresentation analyses were carried out using clusterProfiler version v4.2.2 (Padj cutoff value = 0.05). All statistical analyses were performed using R v4.0.3 (https://www.r-project.org).

### CRISPR Tagging

The EPC1 gene in K562 cells was endogenously tagged with a 3xHA-2xStrep tag. CRISPR-Cas9 strategy with marker-free co-selection using the drug ouabain, as previously described (Agudelo et al. 2017). The guide RNAs (gRNAs) used are described in (Dalvai et al. 2015).

## Author Contributions

D.S. and J.C. designed experiments. D.S., E.G., N. A., and C. L. performed the experiments. C.J.-B., S.B., C. T. and L.H. analyzed the genomic data. D.S. explored the genomic data and rendered the final visualization. M.Q.B. provided patient samples. J.-P.L. performed the mass spectrometry and proteomic analysis. J.-P.L., M.T., A.D., and J.C. supervised and secured the funding. D.S. and J. C. wrote the manuscript.

## Acknowledgements

We thank Valérie Côté, Samarth Thonta Setty, and Ke Xu for crucial technical support. We thank the entire J.C. lab for discussions that improved this study. We thank Professor Nada Jabado for providing EZHIP cDNA and the bmMSC cell lines. We thank Compute Canada for using supercomputers and the Centre Hospitalier Universitaire de Québec-Université Laval proteomic and sequencing platforms. The models were created using the Biorender.

This work was supported by grants from the Canadian Institutes of Health Research (CIHR; FDN-143314, PJT-178367) to J.C., the Government of Québec, the Ministry of Economy and Innovation to B.C., and the Natural Sciences and Engineering Research Council (NSERC) of Canada to J.-P.L. (RGPIN-2017-06124) and the University of Toronto startup funds to M.T. D.S. was supported by Ph.D. scholarship from Fonds de la Recherche Québec-Santé (FRQS) and Fonds de la Recherche Québec-Nature/Technologie (FRQNT). J.-P.L. is a senior FRQS scholar. J.C. held the Canadian Research Chair in Chromatin Biology and Molecular Epigenetics.

## Supplementary figure legends

**Figure S1.**
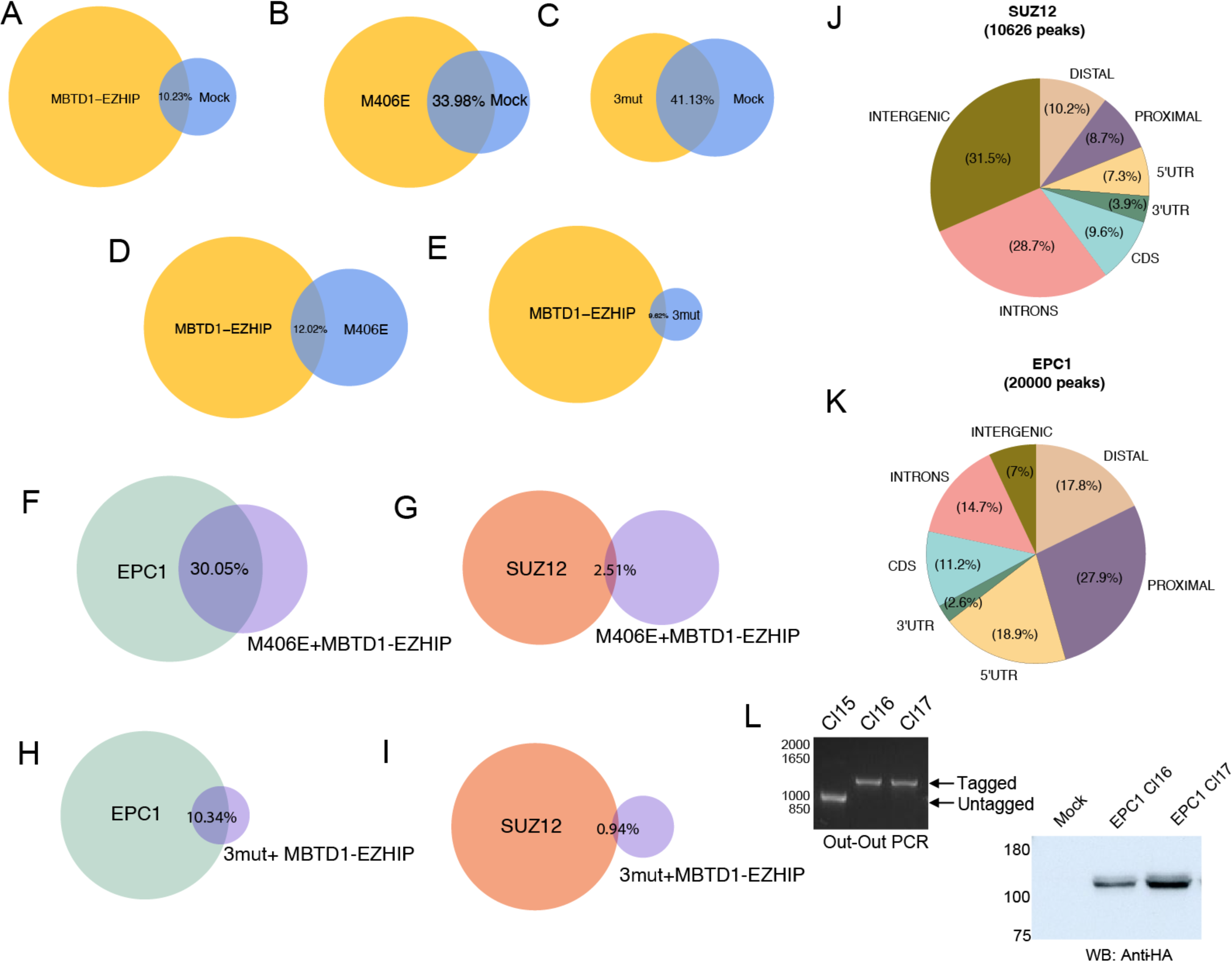
Overlaps of genome-wide location analysis by CUT&RUN of the different fusion proteins with PRC2 (SUZ12) and NUA4/TIP60 (EPC1). **(related to Figure 2) (A-I)** Venn diagrams showing the overlap of the labelled CUT&RUN peaks in K562 cells. **(J, K)** SUZ12 **(J)** and EPC1-HA **(K)** CUT&RUN peaks overlapping with functional elements in the genome**. (L)** Endogenous tagging of EPC1 alleles with a C-terminal 3xHA-2xStrep tag; left panel is out-out PCR to select tagged clones, and the right panel is anti-HA western blot confirmation of the insertion of the tag.

**Figure S2.**
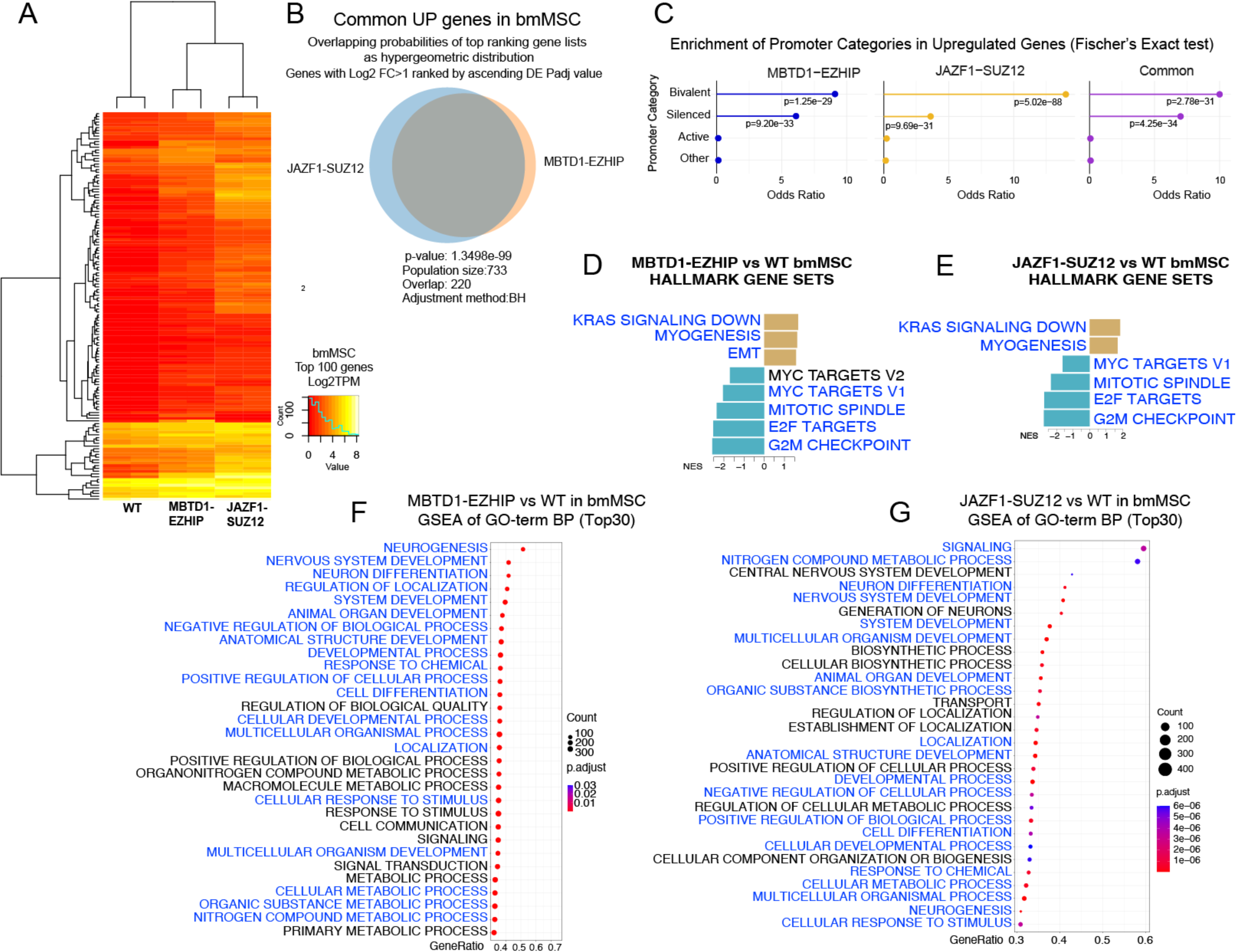
RNA-sequencing followed by differential expression analysis in bone marrow-derived Mesenchymal Stem Cells (bmMSC). **(related to Figure 4) (A)** Hierarchical clustering of the top 100 differentially expressed genes in control compared to bmMSC cells expressing JAZF1-SUZ12 and MBTD1-EZHIP. (B) Venn diagram showing the significant overlap between upregulated genes in JAZF1-SUZ12 and MBTD1-EZHIP expressing bmMSC cells (Log2FC > 1). **(C)** Enrichment of promoter categories in upregulated genes. **(D-E)** Gene Set Enrichment Analysis (GSEA) of differentially expressed genes using the Hallmark-Human Molecular Signatures Database (MSigDB). **(F-G)** Gene Set Enrichment Analysis (GSEA) of differentially expressed genes using the Gene Ontology-Biological Process gene set (GO-BP). Biological processes common between JAZF1-SUZ12 and MBTD1-EZHIP expressing cells are colored in blue.

**Figure S3.**
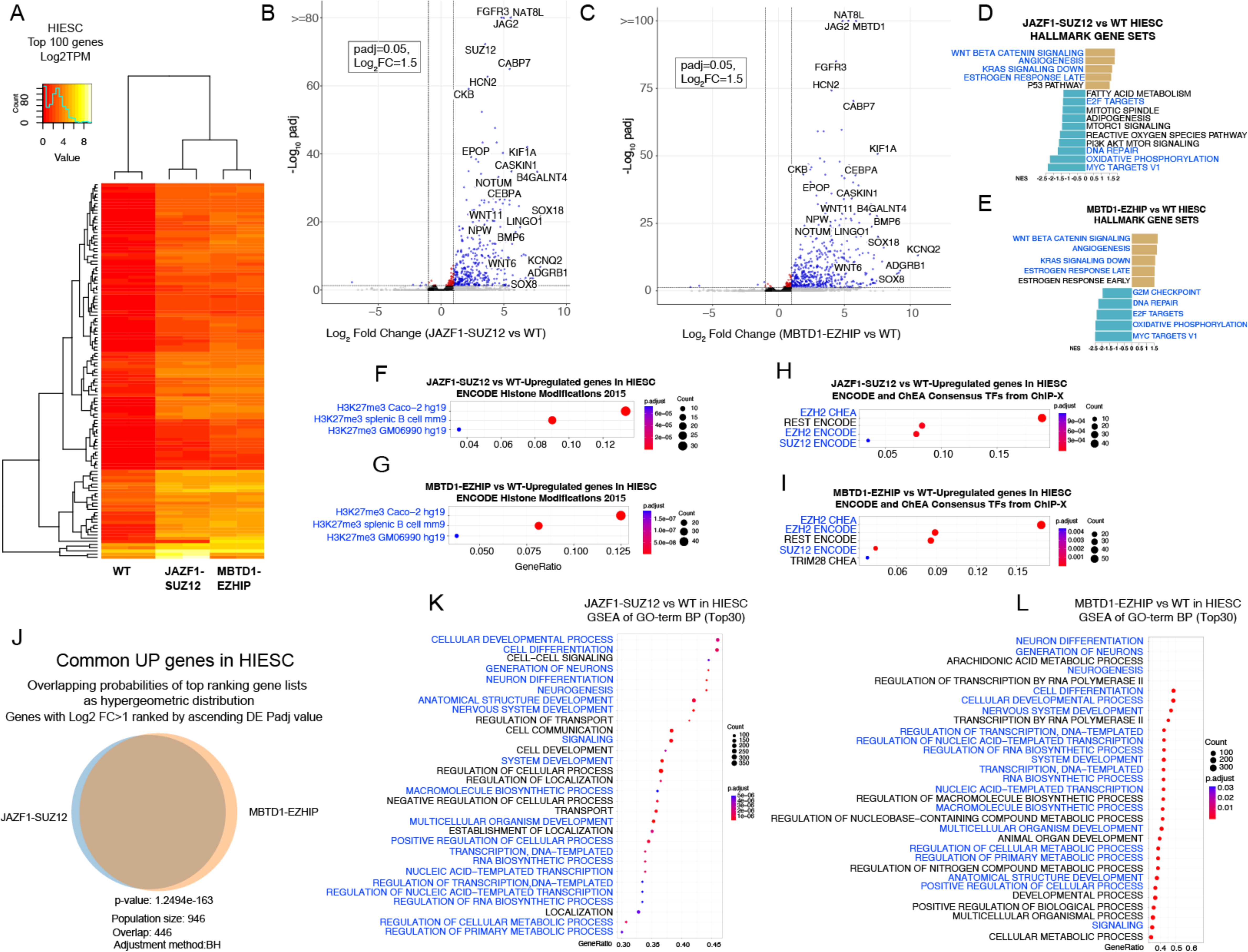
RNA-sequencing followed by differential expression analysis in Human Immortalized Endometrial Stromal Cells (HIESC). **(related to Figure 4) (A)** Hierarchical clustering of the top 100 differentially expressed genes in control compared to HIESC cells expressing JAZF1-SUZ12 and MBTD1-EZHIP. **(B)** RNA-sequencing Volcano plot of HIESC cells expressing MBTD1-EZHIP fusion protein compared to empty vector control/Mock. The graphs show a cut-off at 0.05 for the padj value, and 1.5 for Log2 fold change. An upper limit for -Log10 padj value was set at 80 and 30 at the Y axis, respectively. **(C)** Volcano plot of HIESC cells expressing JAZF1-SUZ12 fusion protein compared to empty vector control/Mock. **(D-E)** Gene Set Enrichment Analysis (GSEA) of differentially expressed genes using the Hallmark-Human Molecular Signatures Database (MSigDB). Gene sets common between JAZF1-SUZ12 and MBTD1-EZHIP expressing cells are colored in blue. **(F-G)** Gene over-representation analysis of upregulated genes using the ENCODE_Histone_Modifications_2015 gene set libraries. **(H-I)** Gene over-representation analysis of upregulated genes using the ENCODE_and_ChEA_Consensus_TFs_from_ChIP-X. **(J)** Venn diagram showing the significant overlap between upregulated genes in JAZF1-SUZ12 and MBTD1-EZHIP expressing HIESC cells. **(K-L)** Gene Set Enrichment Analysis (GSEA) of differentially expressed genes using the Gene Ontology-Biological Process gene set (GO-BP). Biological processes common between JAZF1-SUZ12 and MBTD1-EZHIP expressing cells are colored in blue.

**Figure S4.**
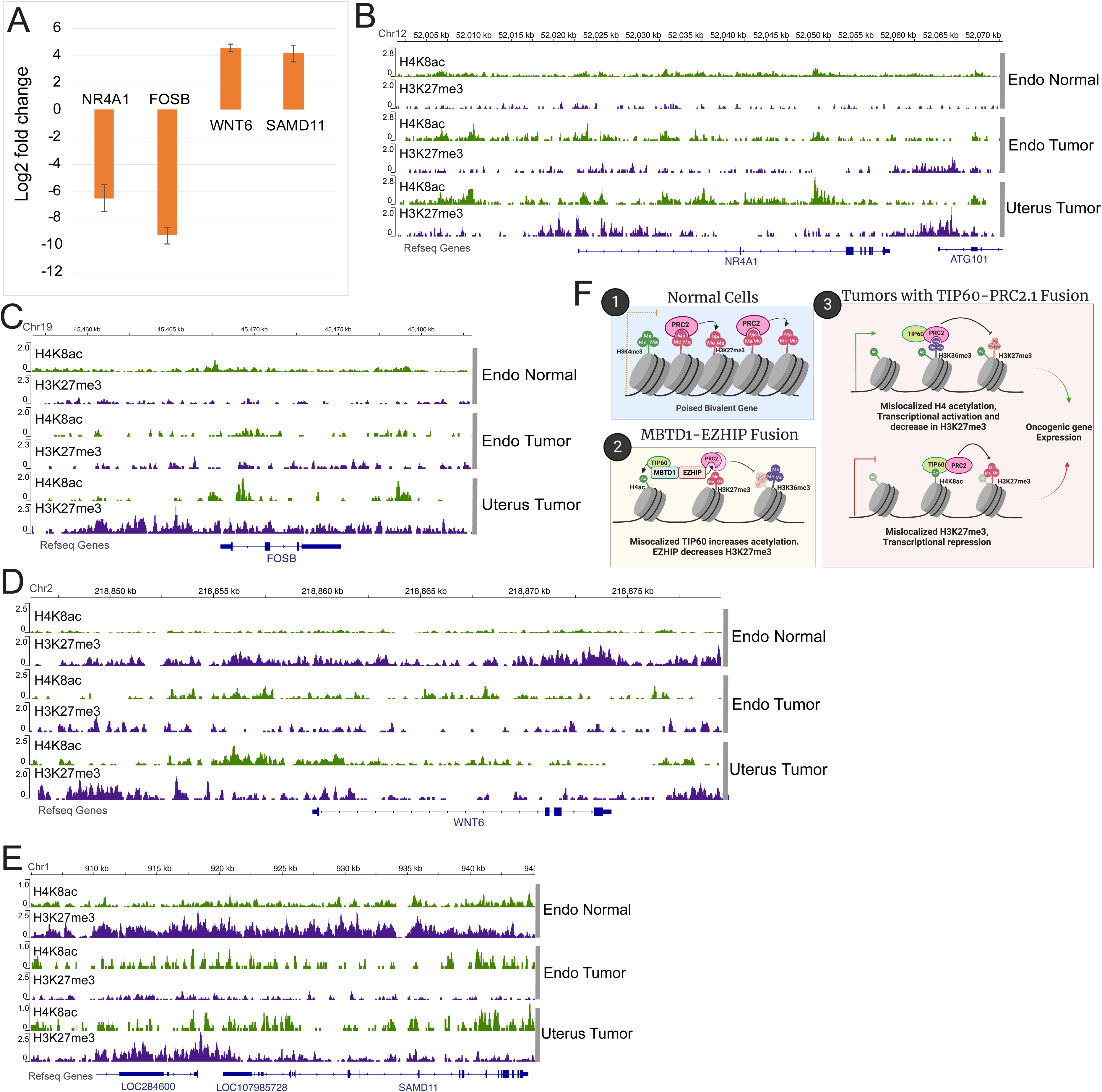
Genome browser views of normalized read counts (RPM) of CUT&RUN-seq at specific genes in patient samples and mechanistic model. **(related to Figure 5) (A)** Selected genes showing change in expression level in JAZF1-SUZ12 expressing patient tumors reported in (Sudarshan et al. 2022) **(B-E)** Representative anti-H4K8ac and anti-H3K27me3 CUT&RUN coverage tracks at differentially expressed genes (shown in A) in normal and patient tumor tissue samples; (B) *NR4A1* (C) *FOSB* (D) *WNT6* (E) *SAMD11.* (F) Model of the molecular mechanism of NuA4/TIP60-PRC2 fusions in LGESS. (1) In normal cells, PRC2 complex occupies repressed or poised chromatin (2) MBTD1-EZHIP fusion protein assembles a chimeric complex combining NuA4/TIP60 and PRC2 complexes. The Fusion-complex decreases H3K27me3 over broad regions through the action or EZHIP and increases H4 acetylation through NuA4/TIP60 activity leading to gene expression changes. (3) Patient tumor samples show conserved mechanism of H4ac increase and H3K27me3 decrease due to the JAZF1-SUZ12 fusion, but also additional mis-localization and increase of H3K27me3 leading to gene repression.

